# Cyclophosphamide chemotherapy induces early p53-directed cytotoxic gene expression changes in ovarian gonadotoxicity

**DOI:** 10.64898/2025.12.04.691500

**Authors:** Xia Hao, Africa Vincelle-Nieto, Arturo Reyes Palomares, Andres Salumets, Armando Reyes-Palomares, Kenny A. Rodriguez-Wallberg

## Abstract

The alkylating chemotherapeutic cyclophosphamide (CPA) is gonadotoxic, commonly resulting in depletion of ovarian primordial follicles and leading to infertility and premature menopause in female cancer patients. However, the mechanisms underlying the early stages of CPA-induced ovarian damage are unclear, limiting our ability to prevent gonadotoxicity. This study provides a comprehensive temporal exploration of the transcriptomic profiles of *ex vivo* intact mouse ovaries exposed to CPA. Analyses of CPA responses from 8 to 36 hours revealed an important early role of p53 signaling. Bioinformatic analyses showed early increases in expression of specific gene clusters associated with apoptosis, DNA damage responses, and cell cycling, while expression of autophagy-associated genes was decreased. Several transcription factors, including E2F family members, likely drive early apoptosis-induced damage and p53 pathway activation. These findings provide mechanistic insight into CPA-induced ovarian toxicity, providing new avenues for the development of protective interventions.

## Introduction

In mammals, females are born with a limited number of ovarian primordial follicles. Most primordial follicles remain dormant during reproductive years but from puberty, waves of primordial follicles are periodically activated to enter folliculogenesis, the developmental process that allows oocyte growth and maturation. However, only a tiny portion of the follicles will be ovulated as a mature egg ready for fertilization during reproductive life, most vanish through follicular atresia. When most of the dormant primordial follicles are exhausted either through activation or atresia during prolonged dormancy, the female becomes infertile. Thus, the size of the pool of primordial follicles determines the female reproductive period^1,2^. Control of dormancy, survival and activation of primordial follicles relies on a complex and interconnected network of molecular signals. Although not yet fully understood, research has shown that this signaling involves crosstalk between granulosa cells (which surround oocytes, providing structural and metabolic support) and oocytes^3^.

Cyclophosphamide (CPA) and other DNA alkylating drugs are used to treat common cancer types in females, such as breast and ovarian cancers. Due to its immunosuppressive effects, CPA is also used in the treatment of autoimmune diseases as well as for pre-conditioning in bone marrow or renal transplantation^4–6^. CPA is a prodrug and its active metabolites interact with DNA to form DNA adducts and double-strand breaks (DSBs), which disrupt DNA replication, leading to cytotoxicity^7^. As a side effect, CPA has strong gonadotoxic effects^8,9^. In female cancer patients, CPA treatment results in ovarian primordial follicle depletion (PFD), which can lead to premature ovarian insufficiency (POI) and infertility.

The mechanisms involved in female gonadotoxicity have been intensively studied but consensus is lacking. Alkylating agents induce DNA damage in cells such as granulosa and oocytes, which is followed by follicular apoptosis in human and rodent models *in vivo* and *in vitro*^10–19^. Consequently, DNA damage-induced apoptosis has been proposed to be the main mechanism for CPA-induced ovarian toxicity^15,20–25^. Indeed, histological studies of *ex vivo* mouse ovaries treated with CPA revealed profound structural changes associated with apoptosis in the ovary after 24 hours of treatment, leading to depletion of the follicle pool^26^. An additional, mutually exclusive, hypothesis suggests that the gonadotoxic effect of CPA is through activation of the PI3K/PTEN/Akt signaling pathway, resulting in over-activation of primordial follicles and thus “burnout” of the ovarian follicle pool^25,27–34^. Although less investigated, there are additional theories including that gonadotoxic chemotherapy damages the ovarian micro-capillary network, which induces cellular stress that results in PFD^11,35–38^. Although most studies support the induction of apoptosis in ovarian follicles in female cancer patients receiving CPA, the precise mechanisms underlying this response are unclear.

In this study, we investigate the temporal dynamics of transcriptomic changes in *ex vivo* prepubertal mouse ovaries treated with CPA. Bulk RNA sequencing of intact ovaries treated with CPA *in vitro* in a time-course up to 36 hours, revealed key roles of p53 signaling, apoptosis and DNA damage responses in the early stages of gonadotoxicity, suggesting that primordial follicles are not reactivated. The identification of these early targets of CPA-induced ovarian damage could reveal targets to protect female fertility during treatment with alkylating chemotherapy in the future.

## Results

### CPA treatment of mouse ovaries in vitro results in transcriptome remodeling and changes in cellular composition

CPA is a prodrug that is metabolized by the cytochrome P450 system, forming 4-hydroxycyclophosphalmide (4-OHC). 4-OHC is then rapidly converted to aldophosphamide, which subsequently produces the active alkylating agents phosphoramide mustard and acrolein^7^. Therefore, studying the effects of CPA *in vitro* requires the use of 4-hydroperoxy cyclophosphamide (4-HC), which generates 4-OHC *in vitro*. To examine the early effects of CPA on ovaries, we performed bulk RNA sequencing (RNA-seq) of intact ovaries from B6CBA/F1 female mice at postnatal day 4 (PND4) cultured in 5 µM of 4-HC^17,26^ for 8, 12, 24 and 36 hours (see Methods). We then compared gene expression profiles at each time point to those of untreated controls to estimate the effects of CPA over time (Fig. 1A, n = 4-5 per group). Ovaries from PND4 mice were chosen as an experimental model because they are mainly composed of primordial follicles, thus allowing the ovarian follicle pool to be captured. Additionally, our histological studies have established that this model system, designed to reflect human-relevant conditions, exhibits primordial follicle depletion in response to CPA^26^.

**Figure 1.**
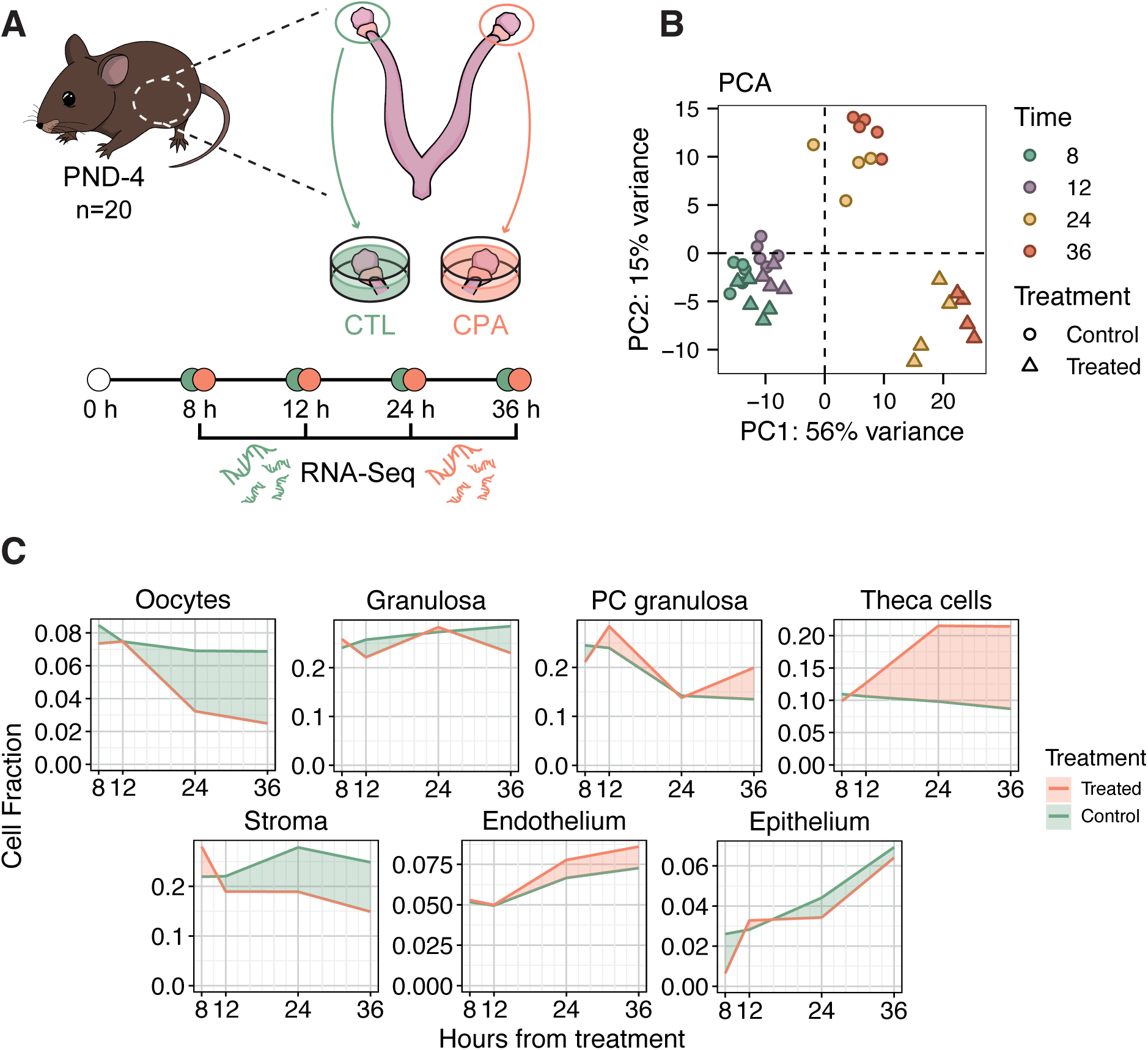
CPA treatment induces major transcriptomic changes in mouse ovaries at 24 h and 36 h. **(A)** Graphical scheme of the experiment. **(B)** Principal component analysis (PCA) using the 3000 most variably expressed genes across samples, showing different time points by color (*8h*-green, *12h*-purple, *24h*-yellow, *36h*-orange), and different conditions by shape (*Control*-circle, *Treated*-triangle). **(C)** Area line plot of the average inferred cellular composition of the ovary per condition across time points. Orange and green indicate a higher fraction of a certain cell type in the CPA-treated and control samples, respectively. PC granulosa, preantral-cumulus granulosa.

RNA-seq data from intact ovaries were homogeneous, and sufficient sequencing coverage was achieved across all samples (Supp. Fig. 1A). In total, 31,936 expressed genes were detected. Principal component analysis (PCA) revealed segregation of samples according to time point and treatment (Fig. 1B). While gene expression variations in response to CPA were detectable at 8 h, specific transcriptomic differences due to treatment were pronounced after 24 h, as visualized in the second principal component (accounting for 15% of the variation) (Fig. 1B). These differences were also noticeable through hierarchical clustering of samples based on their transcriptome and variable genes (Supp. Fig. 1B-C). These results are consistent with histological alterations arising in the ovary after 24 hours of CPA treatment^26^.

To assess the changes in cellular composition caused by CPA, we performed a deconvolution analysis applying CIBERSORTx algorithm^39^ to derive cell type-specific fractions using gene signatures from a single cell RNA-seq dataset of mouse ovary^40^ (Fig. 1C). We observed subtle time-dependent changes in the cellular composition of untreated ovaries in culture (Fig. 1C) but we detected substantial CPA-induced changes in cell types associated with the ovarian follicle (Fig. 1C). In particular, oocytes are significantly depleted to 47% compared with untreated ovaries after 24-hour exposure to CPA (Wilcoxon test p = 0.029) (Supp. Fig. 1D). This decrease aligns with the observed histological depletion of primordial follicles in mouse ovaries after 24 h of CPA treatment^26^. We also found that the proportion of granulosa and preantral-cumulus (PC) granulosa cells showed significant differences between treated and control samples at 36 h: PC granulosa cells were increased in CPA-treated samples (Wilcoxon test p = 0.016), whereas granulosa cells showed a decrease compared to controls (Wilcoxon test p = 0.016) (Fig. 1C, Supp. Fig. 1D). Additionally, the fraction of theca cells (endocrine cells within ovarian follicles) in the treated group was significantly increased from 12 h (Wilcoxon test p = 0.016, p = 0.029 and p = 0.016 at 12 h, 24 h and 36 h, respectively) to more than double the proportion in controls at 24 h and 36 h (2.19 times at 24 h and 2.47 times at 36 h) (Fig. 1C, Supp. Fig. 1D). Although not always significant, changes to the proportion of other ovarian cell types were also observed, such as decreased stromal cells from 12 h onwards in the treated compared to control samples (Wilcoxon test p = 0.19, p = 0.057 and p = 0.016 at 12 h, 24 h and 36 h, respectively) (Fig. S1D), and increased endothelial cells in the treated samples compared with untreated controls at 24 h and 36 h (Wilcoxon test p = 0.057 at 24 h and p = 0.016 at 36 h).

Taken together, our results suggest that CPA treatment alters gene expression and the cellular composition of ovarian tissue, thereby altering the follicular microenvironment, particularly after 24 h of exposure. Nonetheless, more subtle alterations were sometimes detected at earlier time points, suggesting a progressive change in the transcriptomic landscape that becomes more pronounced after 24 h of CPA treatment.

### CPA treatment progressively induces transcriptomic remodeling associated with DNA damage responses and p53 signaling

To identify temporal responses of ovaries to CPA treatment, we performed differential gene expression (DGE) analyses between treated and control samples for each time point (Supp. Table 1). A total of 109, 142, 4410, and 5851 differentially expressed genes (DEGs; 5% false discovery rate [FDR]) were identified at 8, 12, 24 and 36 h, respectively (Fig. 2A). Although subtle expression changes were detected at 8 h, the greater number of DEGs were found at 24 and 36 h (Fig. 2A; Supp. Figs. 2A, 2B, 2C). We identified a core set of 26 common DEGs across all comparisons that exhibited persistent increased gene expression up to 36 h (Fig. 2A, 2B; Supp. Fig. 2A, 2C, 2D). Interestingly, based on CollecTRI database of transcription factor (TF) regulons^41^, 13 of those 26 common DEGs were direct gene targets of p53 (Fisher’s test odds ratio = 35.16 and p = 4.49e-14) (Fig. 2B, highlighted in red). This set of 13 DEGs included genes such as *Cdkn1a* (which encodes p21), and those encoding TFs such as E2F7, both of which are known to have roles in DNA damage responses^42–45^.

**Figure 2.**
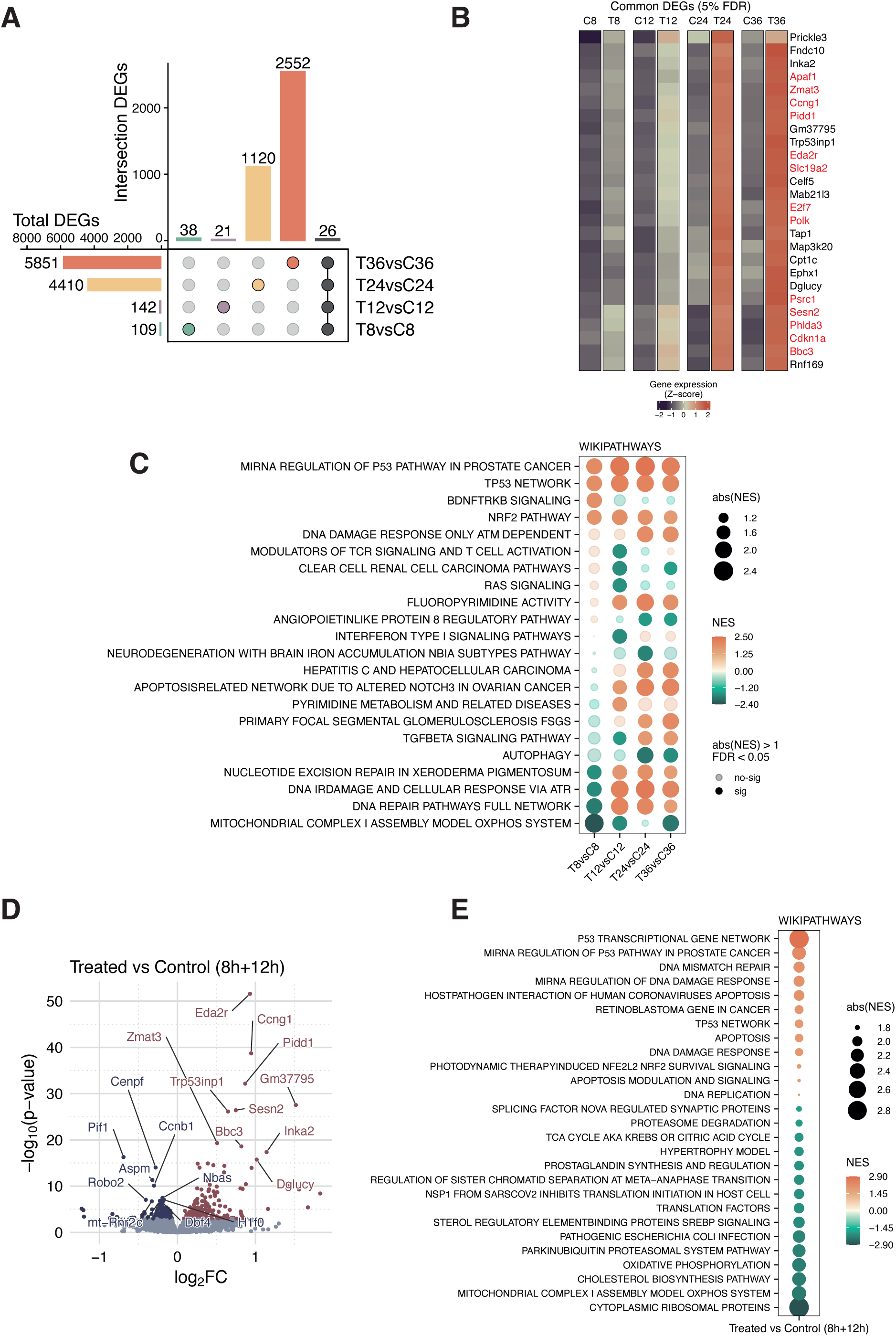
Significant differences in gene expression between CPA-treated and control ovaries arise from as early as 8 h. **(A)** Upset plot of differentially expressed genes (DEGs; adjusted p-value<0.05) across comparisons from the differential gene expression (DGE) analyses. Colored points represent DEGs unique to each comparison, and grey points represent DEGs shared between all comparisons. Left bar plot shows the total number of DEGs per comparison, and top bar plot shows the number of DEGs in each intersection. Only selected intersections are shown (all intersections are represented in Supplementary Fig. 2C). **(B)** Heatmap of mean vst normalized gene expression per group of the 26 common DEGs between conditions across the 4 studied time points. Genes that are transcriptional targets of p53 are highlighted in red (based on CollecTRI regulons). **(C)** Gene set enrichment analysis (GSEA) performed with the statistic values from the DGE analyses for each time point comparison using annotation from Wikipathways database. Top 3 most significant terms with positive normalized enrichment score (NES) and top 3 significant terms with negative NES were selected for representation from each time point comparison, colored in orange and green, respectively. Pathways with 5% false discovery rate (FDR) and absolute NES > 1 were considered significant, non-significant terms are represented with a higher transparency as specified in the legend (no-sig). Dot size represents absolute NES value. **(D)** Volcano plot showing the differential expression levels of genes between treated and control mouse ovary samples from pooled 8h and 12h time points. DEGs (adjusted p-value < 0.05) are represented in dark-red and dark-purple colors if upregulated or downregulated in the treated samples, respectively. Top 10 most significant DEGs (up- and downregulated) are labelled. **(E)** GSEA performed with the statistic values from the DGE analysis pooling 8h and 12h samples, using annotation from Wikipathways database. Top 15 most significant terms with positive NES, and top 15 significant terms with negative NES were selected for representation, colored in orange and green, respectively. Pathways with 5% FDR and absolute NES > 1 were considered significant. Dot size represents absolute NES value.

To further investigate the mechanisms underlying early and progressive CPA responses, we performed a gene set enrichment analysis (GSEA) on the identified DEGs using the WikiPathways database^46^ (Fig. 2C, Supp. Table 2). At 8 h we found a strong enrichment in CPA-treated samples of upregulated genes involved in p53 signaling and other regulatory pathways associated with protective functions against oxidative stress, such as the NRF2 pathway^47^ (Fig. 2C). Interestingly, we also detected a peak enrichment at 8 h of the BDNF/TrkB pathway, which is related to follicle development and oocyte maturation^48^. Conversely, we noted an early and robust downregulation of mitochondrial complex I assembly in CPA-treated samples, consistent with protection from oxidative stress (Fig. 2C). It is noteworthy that DNA damage repair pathways showed a downregulation at 8 h in CPA-treated ovaries, followed by an upregulation from 12 h, while DNA damage response showed an upregulation only from 24 h (Fig. 2C). Over the course of 12-24 h with CPA treatment, we observed upregulation of pathways related to apoptosis and pyrimidine metabolism (Fig. 2C). Conversely, we found that CPA treatment downregulated immune-related mechanisms involving modulators of T-cell receptor (TCR) signaling and T cell activation, interferon type I and transforming growth factor β (TGF-β), the latter being reactivated after 24 h of treatment^49^. Finally, mechanisms associated with autophagy were strongly downregulated at all time points (Fig. 2C).

Because our DGE analysis revealed few but significant changes at early time points (8 and 12 h), we conducted an additional early-onset DGE analysis focused on these time points (Supp. Table 3). To enhance the detection of early alterations in response to CPA, we pooled the 8 h and 12 h samples and compared treated versus control groups, adjusting for CPA exposure time. As a result, we obtained 397 DEGs (5% FDR), where 232 were upregulated and 165 were downregulated (Fig. 2D). All common upregulated DEGs from the individual time point comparisons (Fig. 2B) were found among the top significant upregulated DEGs of this pooled analysis, validating these early changes. By looking at top downregulated DEGs, we found *Pif1*, which encodes a helicase involved in the suppression of apoptosis in human tumor cells^50^. To further inspect the pathways altered in CPA-treated samples compared with controls at these early time points, we again performed a GSEA using WikiPathways database^46^ (Fig. 2E; Supp. Table 4). We found that the strongest signal for upregulation was associated with p53, apoptosis and DNA damage responses (Fig. 2E), which validates our previous observations from the DGE analyses of individual time points (Fig. 2C). Moreover, pathways that were downregulated were associated with translation elements (translation factors, ribosomes), proteosome mechanisms, mitochondrial metabolism (particularly oxidative phosphorylation), and metabolic pathways (TCA cycle, prostaglandin and cholesterol synthesis, sterol signaling) (Fig. 2E).

Overall, these results demonstrate a time-dependent transcriptomic response in ovaries to CPA treatment that specifically upregulates genes associated with p53 signaling and regulation as well as DNA damage and stress response pathways at early time points (8-12 h), identifying initiating molecular processes of gonadotoxicity.

### CPA induces the expression of functional gene clusters associated with apoptosis, DNA damage responses and cell cycle regulation

Our DGE analyses revealed complex time-dependent effects of CPA, with the most pronounced gene expression changes occurring after 24 h of treatment and with fewer but significant changes observed at earlier time points. Thus, we modelled, in a genome-wide approach, the gene expression dynamics using a Partitioning Around Medoids (PAM) clustering strategy to describe the modulation of gene programs represented by the 10,000 most variably expressed genes according to time and in response to CPA (Supp. Table 5). We identified 6 distinct gene clusters (Fig. 3A, 3B), showing small changes starting at 8 h that are strongly enhanced between the 12 and 24 h time points (Fig. 3A, 3B).

**Figure 3.**
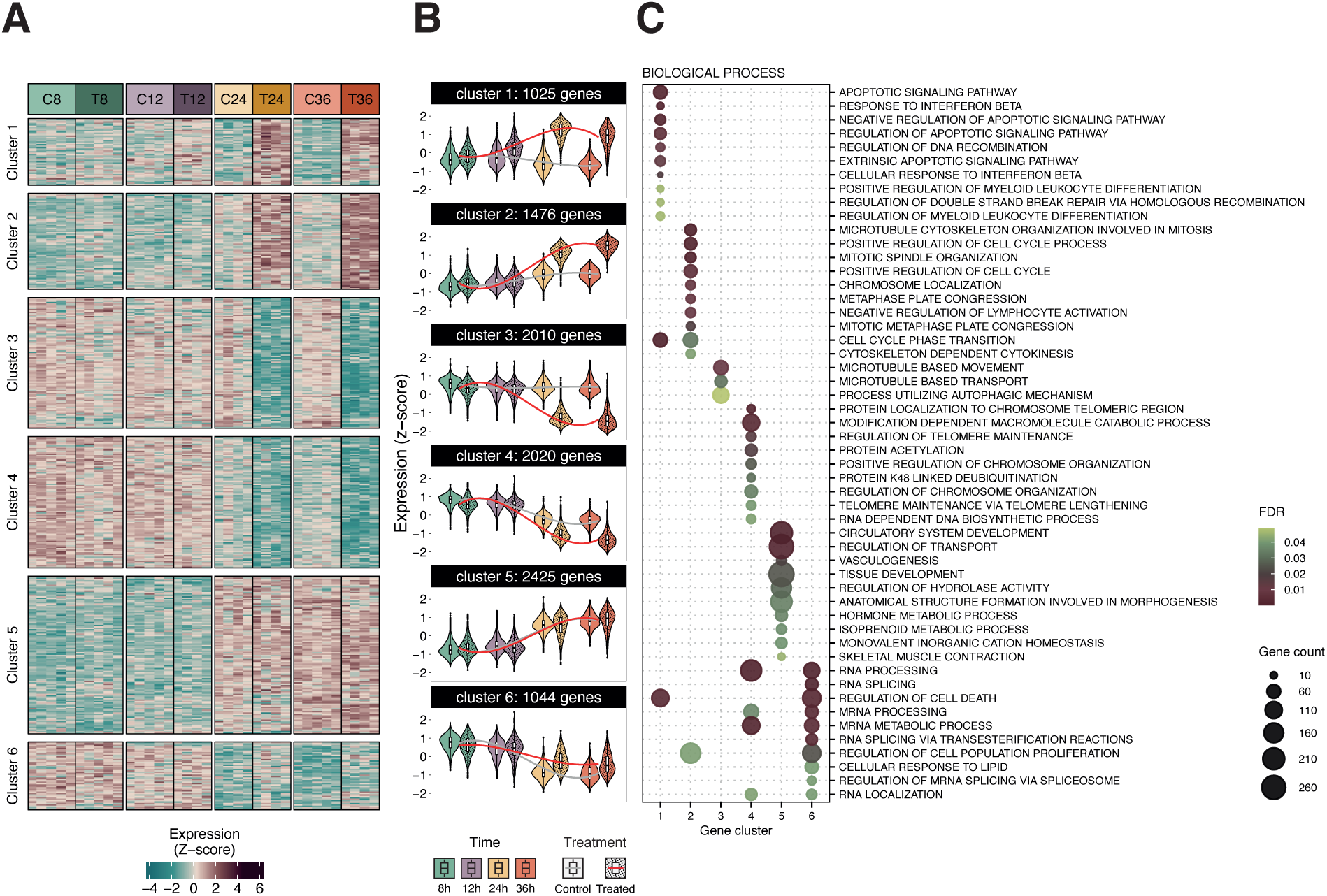
Specific clusters of genes with distinctive expression profiles show incipient changes in the ovaries at 8 h. **(A)** Heatmap of vst normalized gene expression per group showing the 6 clusters obtained after clustering of the top 10,000 most variable genes across samples. **(B)** Violin-box plot of vst normalized gene expression of different gene clusters across groups. Grey (control) and red (CPA-treated) lines represent the expression trends per group across time points. **(C)** Overrepresentation of biological processes from gene ontology (GO) terms for each gene cluster. Top 10 most significant terms were selected from each cluster. Terms with 5% FDR were considered significant. Dot size represents gene count, and color represents FDR.

Overrepresentation analyses of biological processes from Gene Ontology (GO) terms^51,52^ of these 6 distinct clusters revealed specific functions (Fig. 3C; Supp. Table 6). We found three clusters that were associated with CPA treatment, exhibiting both upregulation (cluster 1 and 2) and downregulation (cluster 3) in CPA-treated samples compared with controls. Cluster 1 (comprising 1,025 genes) exhibited pronounced CPA-induced upregulated expression, particularly from 12 h (Fig. 3B). This cluster includes genes associated with apoptosis, cell cycle phase transition, and the regulation of DSB repair via homologous recombination, as well as immune pathways related to the response to interferon β (Fig. 3C). Cluster 2 (1,476 genes) also showed upregulation of gene expression from 12 h after CPA exposure (Fig. 3B). These genes were mostly associated with cell cycling and division, suggesting a possible mechanism in the ovarian tissue to maintain cell proliferation and mitotic activity in the presence of CPA (Fig. 3C). Cluster 3 revealed a robust and sustained downregulated expression of genes associated with microtubule transport and autophagy after 12 h of CPA exposure (Figs. 3B and 3C).

In the remaining clusters (clusters 4, 5 and 6), CPA treatment elicited some differences; however, the observed changes were moderate and appeared largely due to time spent in culture, as gene expression dynamics followed similar trends with and without CPA (Fig. 3A, 3B). Specifically, cluster 4 exhibited a marked downregulation of a core set of genes associated with transcriptional regulation over time, with more substantial effects observed in CPA-treated samples (Fig. 3B, 3C). This cluster includes genes associated with chromosome organization, chromatin modification, and RNA metabolism (Fig. 3C). Lastly, the dynamics of clusters 5 and 6 represented groups of genes with strong alterations in both control and CPA-treated conditions after 24 h of culture, with minor specific CPA contribution (Fig. 3A, 3B). Cluster 5 included upregulation of genes involved in tissue remodeling pathways and cluster 6 showed downregulation of genes related to mRNA processing and RNA localization (Fig. 3C).

These results indicate that the expression of specific gene programs varied in a time-dependent manner after CPA treatment, with some gene clusters responding shortly after exposure. Although some gene expression changes may be attributed to the effects of *in vitro* culture, clusters 1-3 represent biologically relevant responses of ovarian tissue to CPA, characterized by an induction of apoptosis, cell cycle regulation and DNA damage responses while downregulating autophagy, consistent with our earlier results (Fig. 2C).

### The p53 signaling pathway modulates an early response to CPA treatment in ovaries

To deepen our understanding of the temporal responses of ovaries to CPA, we inferred the activity of PROGENy-annotated pathways^53^ implemented in decoupleR^54^ using the expression of their main representative genes. We found a significant increase in p53 related processes as early as 8 h in CPA-treated ovaries compared to controls (Wilcoxon test p = 0.0079 at 8 h) (Fig. 4A, Supp. Fig. 3A). This was the only pathway we found to be significantly upregulated 8 h after CPA, thus further validating our prior findings in this study (Figs. 2B, 2C, 2E). Conversely, PI3K, MAPK and NF-κB were the only pathways significantly downregulated at 8 h (Wilcoxon test p = 0.032) (Fig. 4A, Supp. Fig. 3A). At 12 h of CPA treatment, p53 was also upregulated, together with the JAK-STAT pathway (Wilcoxon test p = 0.016 and 0.032, respectively) (Fig. 4A, Supp. Fig. 3A).

**Figure 4.**
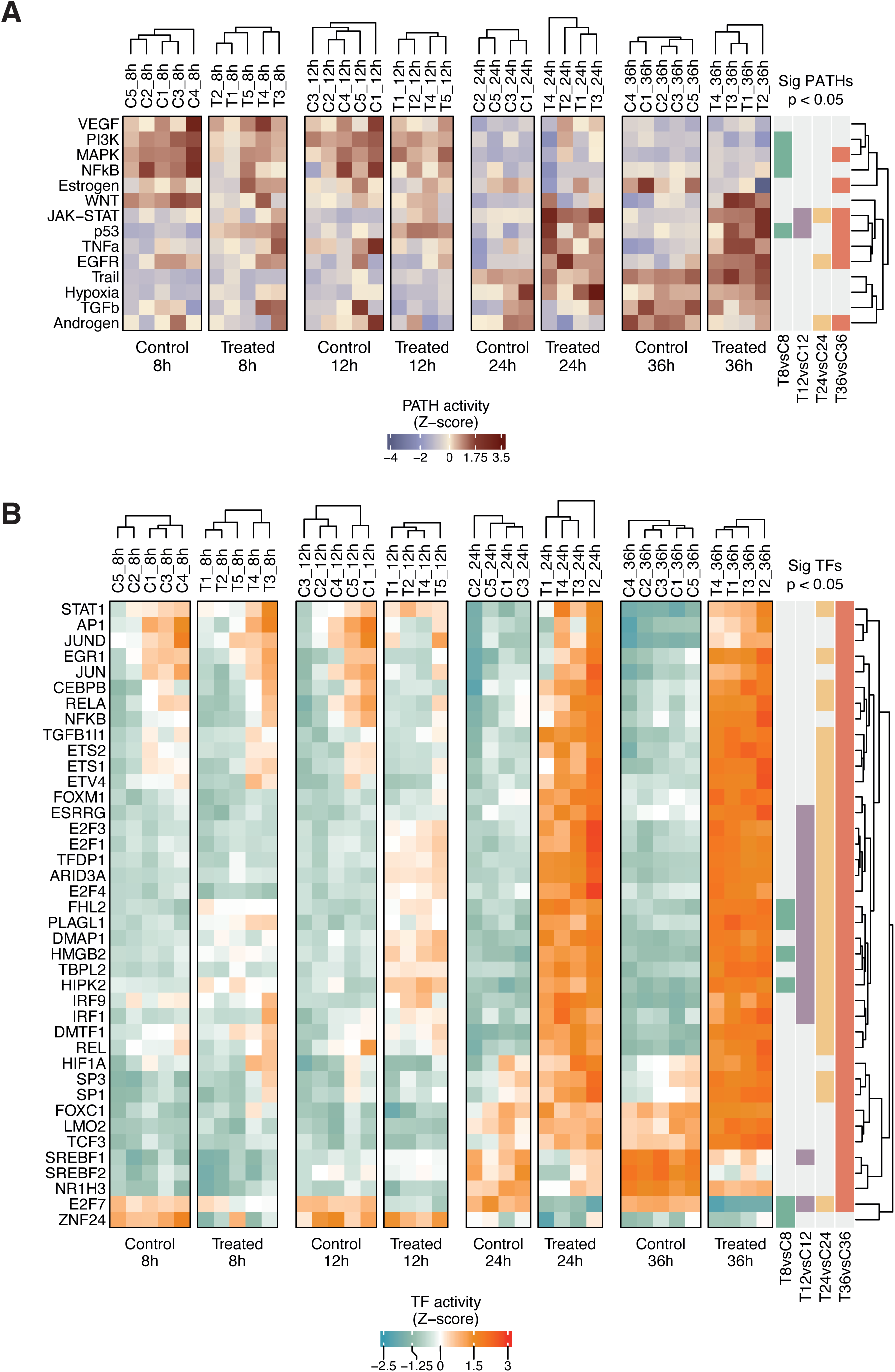
p53 signaling and apoptosis-related TFs show different activities in response to CPA treatment in the ovary. **(A)** Heatmap showing the pathway activity per sample estimated from the expression of the main 500 pathway-related genes retrieved from the PROGENy database. Pathways that have significant activity differences between control and treated samples at each time point are indicated at the right, based on Wilcoxon tests. **(B)** Heatmap showing the TF activity per sample from the 40 most variable TFs across samples. TF activities are estimated from targets’ gene expression using CollecTRI database curated regulons. TFs that have significant activity differences between control and treated samples at each time point are indicated at the right, based on Wilcoxon tests. Human gene symbols are used in this database for clarity, although representation is referring to proteins (TFs). In A and B, CX_Yh represents Sample X in control group at Y hours; TX_Yh represents Sample X in Treated group at Y hours.

CPA-induced changes were also observed in other pathways at later time points. For example, after 24 h, JAK-STAT and EGFR pathways were significantly upregulated (Wilcoxon test p = 0.029) (Fig. 4A, Supp. Fig. 3A), while the androgen pathway was significantly downregulated in the treated samples compared to controls (Wilcoxon test p = 0.029) (Fig. 4A, Supp. Fig. 3A). Finally, after 36 h of CPA treatment, MAPK, JAK-STAT, p53, tumor necrosis factor α (TNFα) and EGFR pathways were significantly upregulated (Wilcoxon test p = 0.032, p = 0.016, p = 0.016, p = 0.016 and p = 0.016, respectively) (Fig. 4A, Supp. Fig. 3A). On the contrary, at this time point estrogen and androgen pathways were significantly downregulated (Wilcoxon test p = 0.032) (Fig. 4A, Supp. Fig. 3A).

There were pathways that exhibited the same changes in both conditions across time points. For example, the activity of hypoxia and TRAIL pathways increased in both groups after 24 h; while PI3K, MAPK and VEGF showed decreased activity in both conditions after 24 h. These changes, as observed in previous analyses (Figs. 3A, 3B), may be caused by *in vitro* culture of *ex vivo* tissue.

We next wanted to identify the modulators involved in the early transcriptional response to CPA treatment, at 8 h and 12 h. For this purpose, we estimated the activities of a curated set of TFs in the CollecTRI database^41^ implemented in decoupleR^54^, which contains an extended annotation of TF regulons, using the expression of their target genes. By examining the 40 TFs whose inferred activity changes the most between samples, we identified different patterns over time (Fig. 4B).

At 8 h, we detected significant changes in the activity of TFs associated with DNA repair, apoptosis and the p53 pathway. In particular, an early significant downregulation of E2F7 activity was observed at 8 h, remaining consistently downregulated along time points in treated samples compared with controls (Wilcoxon test p = 0.016, p = 0.016, p = 0.029 and p = 0.016 at 8 h, 12 h, 24 h and 36 h, respectively) (Fig. 4B, Supp. Fig. 3B). This E2F family member is mainly involved in cell cycle regulation and apoptosis^55^. Interestingly, *E2f7* expression was upregulated across all time points under CPA treatment (Fig. 2B), suggesting an independent regulation at the TF protein level. By contrast, upregulated activities of other E2Fs (E2F1, E2F3 and E2F4) were observed, with higher intensity and significance from 12 h (Fig. 4B, Supp. Fig. 3B).

Interestingly, we noticed a group of 4 TFs that were significantly upregulated from 8 h of treatment across all time points in CPA samples compared to controls: HIPK2, an inducer of apoptosis via activation of p53^56^; HMGB2, a chromatin-associated protein that is involved in mouse ovarian folliculogenesis^57^; PLAGL1, an anti-proliferative zinc finger transcription factor^58,59^; and FHL2, a regulator of cell survival and pro-apoptotic protein^60^, that interacts with both HIPK2 and p53 to induce apoptosis^61^, and is expressed in granulosa cells and also has a role in follicular development^62^ (Wilcoxon test all p < 0.05) (Fig. 4B, Supp. Fig. 3B). A less consistent early downregulation was observed for ZNF24, a zinc finger transcriptional repressor involved in diverse biological processes such as angiogenesis^63,64^. This is the only TF that showed significant changes exclusively at 8 h (Wilcoxon test p = 0.032) (Fig. 4B, Supp. Fig. 3B).

Taken together, these findings indicate a dynamic reconfiguration of transcriptional programs over time following CPA exposure, characterized by TFs related to apoptosis, DNA repair and follicular development, highlighting the coordinated regulatory mechanisms underlying the early ovarian response to treatment.

Examination of the TF regulons revealed that p53 regulates the expression of E2F7, HIPK2, PLAGL1 and FHL2, while HIPK2 and PLAGL1, in turn, also regulate p53 expression. Since this early response of TFs pointed to a major activation of p53 signaling, we built a regulatory network centered on p53 using its targets from the 26 common DEGs that we found significantly associated with CPA treatment across all time points (Fig. 5A). We also included in the network the TFs that significantly changed across all time points that were directly regulated by p53 (E2F7, HIPK2, PLAGL1, FHL2), and added the rest of the E2F family mentioned in the results above (E2F1, E2F3, E2F4) to explore in detail this family of TFs. When observing the network, 3 genes were predominantly regulated by multiple TFs: *Cdkn1a*, *Bbc3* and *Apaf1*, suggesting that these are key drivers of the early response to CPA. Altogether, these observations suggest that p53 is a key modulator of the early ovarian response to CPA treatment. As p53 signaling induces apoptosis, this likely directs the gonadotoxic effect of CPA on ovarian tissue.

**Figure 5.**
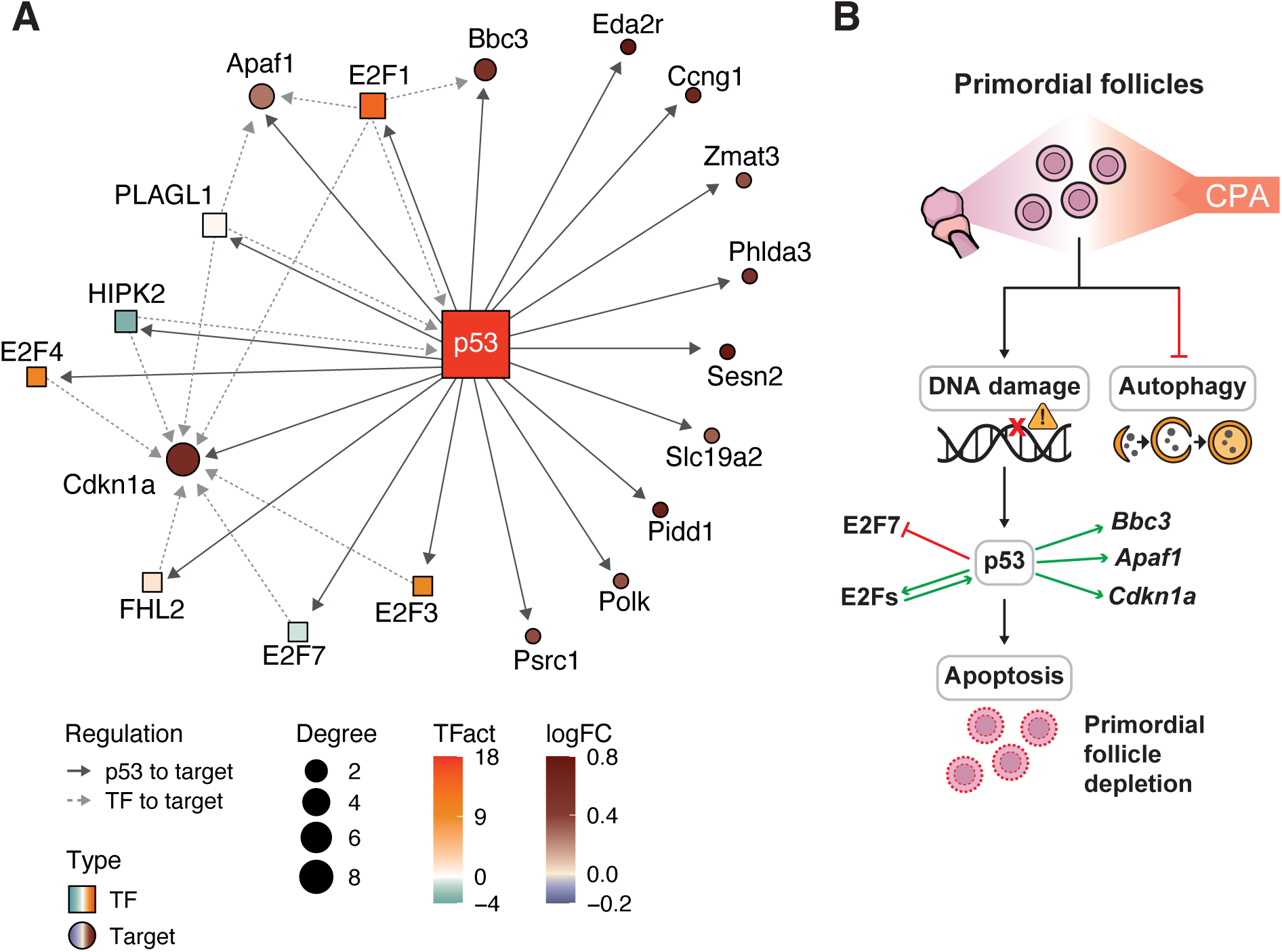
Regulatory network elucidates a potential modulation of apoptosis by p53 signaling. **(A)** Network of p53 target genes that are differentially expressed between conditions across all time points, p53 target TFs with differential activity between conditions from 8 h across all time points (E2F7, HIPK2, FHL2, PLAGL1), and p53 target E2Fs from the TF activity analyses (E3F1, E2F3, E2F4). Target genes have been established based on CollecTRI database curated regulons. TFs are represented with a square and target genes with a circle. Color of target genes represents fold change between treated and control samples at 8 h, color of TFs represents mean TF activity at 8 h in treated samples. **(B)** Schematic overview of the proposed molecular pathways mediating the CPA-induced effects identified in this study.

## Discussion

This study explored the transcriptomic dynamics in *ex vivo* PND4 mouse ovaries for up to 36 h after CPA exposure, aiming to reveal the mechanisms that drive CPA-induced primordial follicle depletion. Notably, the impact of CPA on transcriptional programs manifested early (after 8 h), pointing to p53-mediated DNA damage-driven apoptosis as an early response to CPA, rather than activation of dormant primordial follicles. Transcriptional profiles of specific gene clusters further highlighted DNA damage, apoptosis and autophagy as the main molecular pathways that are altered shortly after exposure. Moreover, some TFs showed significant differential activity between conditions from 8 h onward, suggesting their potential role as drivers of the observed responses through interactions with p53 and regulation of key differentially expressed target genes across all time points.

Our analyses revealed that the most pronounced transcriptional response caused by CPA treatment is triggered at 24 h. At this time point, cell type deconvolution revealed depletion of almost half of the oocytes in the CPA-exposed ovaries, which is consistent with the observed histological depletion of primordial follicles induced by CPA^26^. Conversely, non-germ cells such as theca cells displayed CPA-associated increases, which were detectable as early as 12 h after treatment. Granulosa and PC granulosa cells showed more stable profiles across the experiment, although at 36 h of CPA treatment, the fraction of granulosa cells decreased, which could be attributed to the disruption of follicle structure, either due to oocyte loss or depletion of neighboring follicles^26^. The increase in theca cells is quite interesting. Theca cells are present in growing follicles, indicating the secondary stage of folliculogenesis^65^, and they collaborate with granulosa cells to support follicular growth^66^. Their increase in response to CPA may reflect an as-yet unidentified protective mechanism that supports follicular recovery after gonadotoxic damage, or alternatively may contribute to the previously suggested chemotherapy-induced overactivation of follicles^67^. Nonetheless, further studies are needed to clarify why the proportion of theca cells increases following CPA exposure. In the context of follicular overactivation, the observed upregulation of genes related to cell cycling and division could further support this theory. However, the downregulation of the PI3K pathway is notable, given its role within PI3K/PTEN/Akt signaling axis that drives primordial follicle overactivation^25,27–34^, finally suggesting that this overactivation process is not occurring.

Across all time points, CPA treatment led to significant increases in the expression of a specific set of 26 genes, which were strongly associated with p53 signaling. Moreover, clustering analyses revealed distinct time- and treatment-specific gene expression patterns, showing a gradual upregulation of apoptosis-associated genes and a progressive downregulation of autophagy-associated genes. Furthermore, analyses of TF activity demonstrated that CPA exposure induced significant changes in TFs related to the activation of p53 signaling and apoptosis, including several members of the E2F family^55^. Together, our data indicate that a DNA damage-response transcriptional regulatory network that activates p53 signaling is induced as early as 8 h of CPA exposure.

The interplay between p53 and the E2F family suggests that the E2F-mediated entry into S-phase can, under conditions of deregulated or excessive E2F activity, trigger p53-dependent apoptosis^55^. In our study, we used doses of CPA that are able to induce apoptosis *in vitro*, applied within an experimental model developed to reflect conditions relevant to human exposure, as previously reported^17,26^, and we observed an early downregulation of E2F7 activity from 8 h of CPA treatment and a subsequent upregulation of E2F1 and p53 signaling that ultimately reflect an apoptotic transcriptional program^68^. In this context, we found DEGs associated with p53 and the E2F family: *Bbc3*, a pro-apoptotic member of the BCL-2 family^69^, and *Apaf1*, which is involved in the activation of apoptosis in granulosa cells^70^. Indeed, our regulatory network analysis highlighted the connection between p53 and its transcriptional targets E2Fs and *Cdkn1a* (which encodes p21) in the potential modulation of the cell cycle and apoptosis^71^. Notably, *E2f7* expression increases from 8 h onward, even as its TF activity declines, suggesting a feedback mechanism to compensate for its reduced functional activity at the TF protein level. CPA induces a variety of DNA damage lesions, including intra- and interstrand cross-links and DBSs^67^, which could activate homology-directed repair (HDR) mechanisms in dividing cells. E2F7 mediates p53-dependent cell cycle arrest in response to DNA damage^44^, but has also been linked to the negative regulation of HDR machinery^45^. Consistent with this, we observed a depletion of E2F7 activity accompanied by an increased expression of genes involved in HDR (*Bard1*, *Rad51*, *Rbbp8*-CTIP)^45^, suggesting the induction of HDR pathways in response to CPA. Since HDR is associated with dividing cells, whereas oocytes in primordial follicles are dormant and non-cycling, we cannot exclude that proliferative granulosa precursors present in early postnatal ovaries could contribute to this signal. Nevertheless, accumulating evidence indicates that oocytes from primordial follicles, most of them in meiotic prophase arrest which corresponds to the G2 phase of the cell cycle, preserve a robust HDR capacity, in particular functional homologous recombination (HR) mechanisms, to maintain long-term genomic stability^72–74^. This supports the idea that oocytes may also engage these pathways in response to CPA-induced DNA damage, although if the extent of lesions exceeds their repair capacity, the resulting persistent DNA damage response can lead to apoptosis.

In parallel, we observed downregulated expression of genes associated with autophagy across all time points, supporting previous histological observations^26^. These changes were accompanied by upregulated expression of genes associated with the NRF2 pathway (related to autophagy processes via p62^75^) which, together with Keap1 and ARE, establish an antioxidant pathway against oxidative stress and DNA damage^76^. A recently identified mechanism can upregulate NRF2 when autophagy is inhibited in diverse cancer cell lines^77^, which also seems to be reflected in our data.

We detected a strong early-onset (8 - 12 h) downregulation of the mitochondrial complex I assembly pathway after CPA exposure. This is a recently characterized and evolutionarily conserved oocyte-specific mechanism to prevent the accumulation of reactive oxygen species (ROS) and resulting damage, which preserves homeostasis in this long-lived cell type^78^. Indeed, ovarian tissue is also known to have distinct proteostasis mechanisms that allow increased protein longevity^79^, and in our pooled DGE analysis of 8 + 12 h we observed a significant downregulation of proteasome mechanisms. Together, these findings imply that CPA exposure could be targeting two core mechanisms linked to the maintenance of healthy ovary longevity, possibly contributing to oocyte toxicity. Surprisingly, after 12 h of CPA treatment the pathway activity analyses also showed upregulation of JAK-STAT signaling, which plays roles in ovarian reserve maintenance, follicle activation and development^80–82^.

While our findings reveal early transcriptional responses to CPA in as *ex vivo* culture model using neonatal mouse ovary, the use of this mouse model may not fully reflect human ovarian physiology. In addition, due to the limitations of bulk RNA-seq, single-cell resolution approaches and functional assays in future studies will be crucial to further address these findings. However, our model encompasses the critical window during which ovarian damage is initiated, enabling the characterization of the specific molecular events underlying gonadotoxicity at very early stages, before evident tissue damage becomes apparent. In addition, optimization of intact whole-ovary culture preserved the histological structure and maintained cell-cell interactions, which are essential for the follicular microenvironment and oocyte function.

Together, our results describe the temporal transcriptional response of ovaries to CPA exposure, revealing multiple axes that could potentially be explored as therapeutic targets to prevent follicle loss in the application of fertility preservation strategies in cancer patients. Our findings point to specific potential targets, including apoptotic inhibitors, in line with previous proof-of-concept experimental studies^14^, while underscoring the challenges of broadly applying apoptosis inhibitors in cancer patients. In particular, components of the p53 pathway, like *Cdkn1a*, emerge as early responders to CPA and may represent more selective candidates for ovarian protection.

## Materials and Methods

### Animals and ovarian tissue retrieval

Inhouse bred B6CBA/F1 female mice PND4 (n=20) were used for the experiments. The mice were sacrificed and ovaries were collected in Leibovitz 15 medium containing 10% fetal bovine serum (FBS), penicillin (100 IU/mL) and streptomycin (100 µg/mL). Intact ovaries were exposed by puncturing the ovarian bursal with Micro-Fine U-100 insulin syringes (0.3 mL, BD Medical) under a stereomicroscope (Nikon^®^). This experiment was approved by Karolinska Institutet and the regional ethics committee for animal research in accordance with the Animal Protection Law, the Animal Protection Regulation, the Regulation of the Swedish National Board for Laboratory Animals, identified as Dnr 1372 (Date 2018-01-24).

### Grouping and 4-HC treatment

Intact ovaries were allocated to either 4-HC group (n=20) or control group (n=20), ensuring ovaries of the same mice were allocated into different groups. For culture, ovaries were evenly distributed on top of the membrane’s surface of the Millicell^®^ cell culture inserts (maximum 5 ovaries per insert) soaked with 250 µL of culture medium containing 4-HC solution (Santa Cruz, SC-206885) or an equal amount of the solvents of 4-HC (control group). 4-HC was dissolved in anhydrous tetrahydrofuran and dimethyl sulfoxide and freshly added to culture medium before use, the final content of each solvent was 0.03%, v/v. The culture medium was α-minimal essential medium GlutaMAX supplemented with 10% FBS, transferrin (25 µg/mL), penicillin (5 IU/mL) and streptomycin (5 µg/mL). Culture medium and inserts were pre-equilibrated in a humidified incubator at 37°C and 5% CO_2_ in air. The final concentration of 4-HC was 5 µM, which was selected based on our previous study^26^ and referring to a study that reported 4-HC could induce damage to primordial and small primary follicles at this dose^17^. Incubation was performed under humidified conditions (95%) at 37°C and 5% CO_2_. At 8, 12, 24, 36 hours after the initiation of culture, 5 ovaries from each group were individually collected for snap-freezing by direct immersion in liquid nitrogen and stored at −80°C till further processing. Samples were labeled to indicate, in this order, the experimental condition (C: Control, T: Treated), the replicate number within the group (1 to 5), and the time point (8h, 12h, 24h, 36h): for example, C1_8h: Control sample 1 at 8 h. Sample groups were labeled to indicate, in this order, the experimental condition, and the time point: C8: Control at 8 h, C12: Control at 12 h, C24: Control at 24 h, C36: Control at 36 h, T8: Treated at 8 h, T12: Treated at 12 h, T24: Treated at 24 h, T36: Treated at 36 h, five replicates in each.

### RNA extraction and sample quality control

Total RNA from each ovary was individually extracted using QIAzol lysis reagent and further processed with the miRNeasy Micro Kit (QIAGEN, Germany) following the manufacturer’s instruction. The RNA quantity was detected by Qubit^TM^ 3 Fluorometer (Invitrogen^TM^) and RNA purity and integrity were evaluated by 2200 TapeStation (Agilent Technologies). Only qualified samples (RNA Integrity Number > 6.5, total RNA > 1 μg) were used for following steps, which were performed by Novogene Co., Ltd, UK.

### RNA library preparation

Total RNA was extracted from each ovary sample using QIAzol Lysis Reagent and purified with the miRNeasy Micro Kit (QIAGEN, Germany), following the manufacturer’s instructions. RNA yield and integrity were assessed using a Qubit™ 3 Fluorometer (Invitrogen™) and a 2200 TapeStation (Agilent Technologies), respectively. Only samples with RNA Integrity Number > 6.5 and total RNA > 1 μg were used. Messenger RNA was isolated from total RNA using poly(T) oligo-conjugated magnetic beads. After fragmentation, first-strand cDNA synthesis was performed using random hexamer primers, followed by second-strand synthesis. cDNA libraries were then constructed through end repair, A-tailing, adapter ligation, size selection, PCR amplification for 14-18 cycles, and purification. Final cDNA insert size was 250-300bp. Libraries were sequenced on an Illumina NovaSeq 6000 platform, using paired-end 150 bp reads.

### RNA-seq data processing

Quality control was performed using FastQC software v0.11.9^83^, and those samples that passed quality control were used for alignment. Paired-end reads were mapped to mouse genome assembly (GRCm39) provided by the GENCODE database (vM29 - ENSEMBL release 106) using STAR v2.7.9a^84^ and reads which were uniquely mapped and in correct pairs were retained. Gene-level abundances were estimated using the gene counts option from STAR software for GENCODE GRCm39 vM29 annotation using the primary assembly annotation gtf file. Downstream analyses were performed using R v4.3.2^85^ and Bioconductor v3.18^86^ available packages.

### RNA-seq data analysis

**Quality control.** Quality control of sample similarity between conditions (CPA-treated versus Control) and time points (8 h, 12 h, 24 h, 36 h) was based on principal component analysis (PCA) and hierarchical clustering on transformed gene counts (variance stabilized). Previously, low expressed genes (i.e. genes with a mean expression of < 10 reads across all samples) were removed from further downstream analysis, keeping 31,936 genes. PCA was performed with the top 3,000 variable genes using the DESeq2 package v1.42.1^87^. This procedure allows us to assess any batch effect prior differential expression analysis. Following this quality control step, four ovary samples were detected as outliers and removed from the analyses (Sample 4 of C24, C4_24h; Sample 3 of T12, T3_12h; Sample 5 of T24, T5_24h; Sample 5 of T36, T5_36h).

**Cell deconvolution analysis.** CIBERSORTx tool^39^ was used to estimate the relative fraction of the different ovarian cell-types in each sample. For this purpose, the gene count matrix was CPM-transformed and used as the mixture matrix. Signature matrix was obtained from single cell RNA-seq data published in Niu and Spradling, 2020^40^, retrieved from GEO database supplementary files under accession number GSM4643738 (postnatal ovary from developmental stage P5). Single cell data was preprocessed and analyzed using Seurat R package v4.3.0^88^. Cell populations were established using ovarian mouse marker genes described in Niu and Spradling, 2020^40^ and ovarian human marker genes described in Wagner et al., 2020^89^. ENSEMBL gene IDs were converted to gene symbols using GENCODE vM29 annotation. Finally, both matrices were submitted to the tool for imputing the cell fractions, specifying B-mode batch correction, 100 permutations and absolute mode. Cell types with estimated cell fractions below 1% were considered undetected.

**Wilcoxon Test.** The Wilcoxon test to evaluate p-values on graphs were performed using the ggsignif R package v0.6.4^90^.

**Differential gene expression analysis.** Differential gene expression analysis (DGE) of gene counts was carried out for ovarian samples using the DESeq2 package v1.42.1^87^ with default parameters and using the following model: *“∼ Group”* to contrast treated versus control samples independently for each of the 4 time points. Genes with an adjusted p-value < 0.05 were considered differentially expressed genes (DEGs), unless stated otherwise. For the analysis of pooled 8 h + 12 h samples, the same parameters were used using the model *“∼ Time + Treatment”* to contrast treated versus control samples, and correcting by time point to get only differences associated with the CPA-treatment condition.

Upset plots were performed using the R package ComplexUpset v1.3.3^91^. Heatmaps were performed using ComplexHeatmap R package v2.18.0^92^.

**Gene set enrichment analysis.** Gene Set Enrichment Analysis (GSEA) was performed with the clusterProfiler R package v4.10.1^93^ using the statistic measure from the DGE analysis for each contrast. The gene sets were established by WikiPathways^46^ downloaded from Molecular Signatures Database (MSigDB) for mouse data^94,95^ using the msigdbr R package v7.5.1^96^.

**Gene clustering.** The clustering of genes was performed based on the partitioning around medoids (PAM) algorithm using the cluster R package v2.1.6^97^, across 8 different sample groups (Control_8h, Treated_8h, Control_12h, Treated_12h, Control_24h, Treated_24h, Control_36h, Treated_36h) using mean gene expression per group. The number of clusters (k) was selected based on average silhouette values and downstream exploration via gene set overrepresentation analyses.

**Gene ontology overrepresentation analysis.** Overrepresentation of Gene Ontology (GO) terms for biological processes was performed using the clusterProfiler v4.10.1^93^ using the compareCluster() function and the Gene Ontology database^51,52^, downloaded from Molecular Signatures Database (MSigDB) for mouse data^94,95^ using the msigdbr R package v7.5.1^96^. Only expressed genes were used as background.

**Pathway activity inference analysis.** The activities of 14 different pathways, retrieved from the PROGENy database^53^, were assessed using the gene expression of their top 500 genes involved, by implementing ULM through the decoupleR R package v2.8.0^54^.

**Transcription factor activity inference analysis.** The regulatory activities of TFs were estimated per sample from gene expression data by implementing a Univariate Linear Model (ULM) in TF-target regulons from the mouse CollecTRI gene regulatory network^41^ using the decoupleR R package v2.8.0^54^.

Activities were used to calculate the standard deviation of each TF across samples. Ultimately, we utilized standard deviation as a metric to evaluate variation in TFs, we ranked them, and selected 40 TFs with the highest values.

**Gene regulatory network.** The gene regulatory network depicted p53 target genes differentially expressed between conditions at the 4 measured time points, p53 target TFs with differential activity between conditions from 8 h across all time points (E2F7, HIPK2, FHL2, PLAGL1), and p53 target E2Fs from the TF activity analyses (E3F1, E2F3, E2F4), based on curated regulons from the CollecTRI database^41^. The network was created using the igraph R package v2.0.3^98^.

## Acknowledgements

We would like to acknowledge all the supports from the Preclinical Laboratory at the Karolinska University Hospital, Huddinge, Sweden and the Department of Oncology and Pathology, Karolinska Institutet for providing the excellent platform for research. We would also like to acknowledge the computing resources and technical support provided by the SCBI (Supercomputing and Bioinformatics) Center of the University of Málaga in the Red Española de Supercomputación (RES).

## Author Contributions

X.H., Art.R.P., Arm.R.P. and K.A.R.W. contributed to the conception and design of the study. X.H. performed experimental work. A.V.N. carried out bioinformatic analyses and prepared the figures under supervision of Arm.R.P. X.H. and A.V.N. wrote the initial draft of the manuscript. All authors contributed to draft discussion, manuscript review and revisions. Arm.R.P and K.A.R.W. supervised the study, provided project administration and funding. All authors contributed to the preparation of the paper and approved the submitted version.

## Funding

This research was supported by the Swedish Childhood Cancer Foundation (PR2016-0115, PR2020-0136), the Swedish Cancer Society (CAN 2017/704, 20 0170 F), the Swedish Research Council (Dnr 2020-02230), the Swedish Research Council Collaboration Research Program (NSFC-VR 8211101255), Radiumhemmets Research Funds Grant for clinical researchers (2020-2025), the Stockholm County Council (FoUI-953912), and the Karolinska Institutet Research grants in pediatrics from the Birgitta and Carl-Axel Rydbeck Donation (2020-00339) to K.A.R.W. X.H. was supported by the Estonian Research Council (KOHTO14). A.V.N. was supported by the Regional Grant for Predoctoral Research FPI-CAM from the Regional Government of Madrid (A281-CT65/22). Art.R.P. was supported by Beatriz Galindo Program from the Spanish Ministry of Science, Innovation and Universities (BG23/00015), and the Spanish Ministry of Science, Innovation and Universities with further support from FEDER and UE (PID2024-160756OA-I00), and the Karolinska Institutet Research grants (2024-02566).

## Conflict of Interest

The authors declare that they have no competing interests, and the research was conducted in the absence of any commercial or financial relationships that could be construed as a potential conflict of interest.

## Data Availability

The data of this study will be made publicly available in the ENA Nucleotide Archive repository upon publication.

## SUPPLEMENTARY FIGURES

**Supplementary Figure 1.**
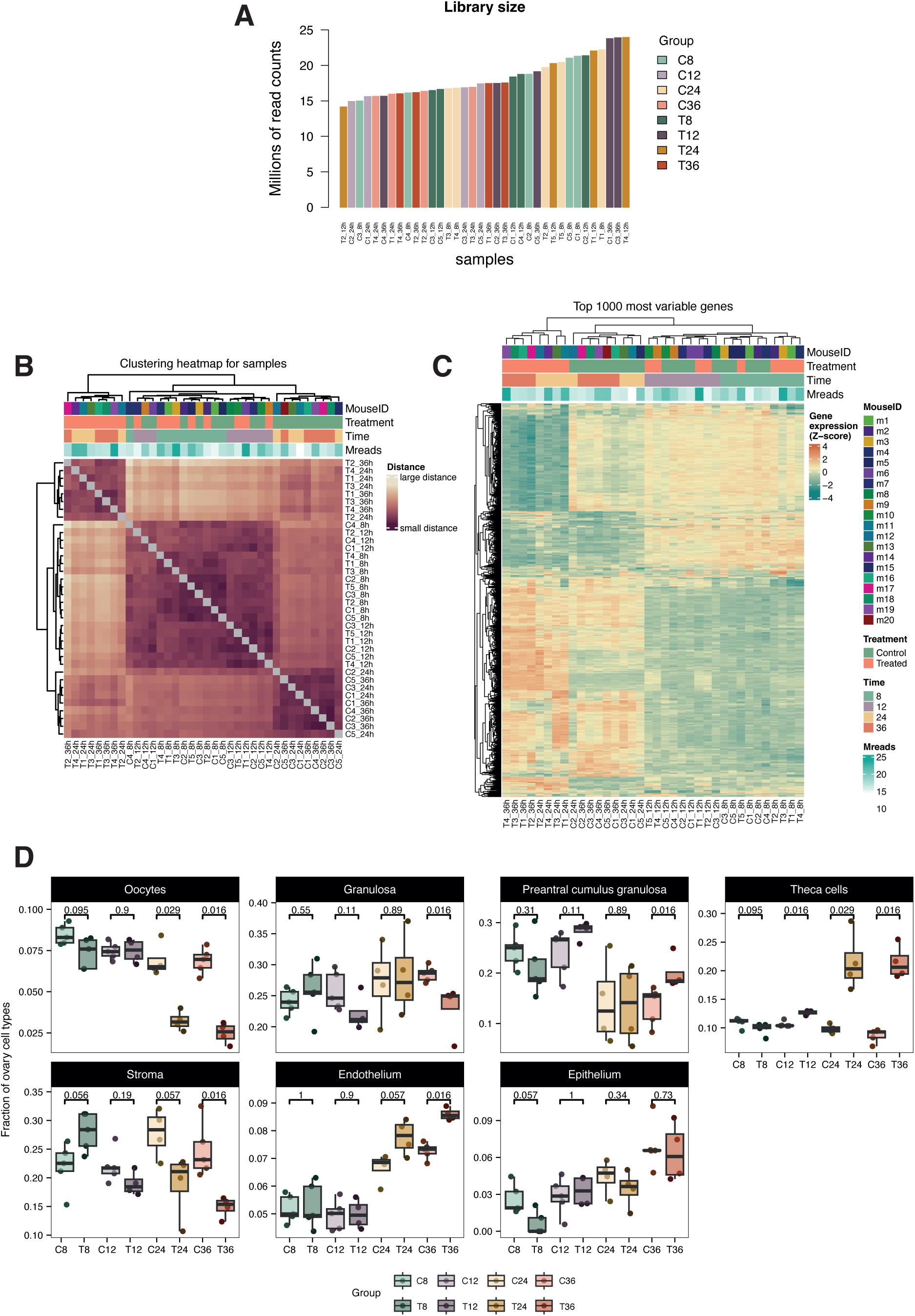
Exploratory data analysis of profiled ovarian mouse samples and cell deconvolution. **(A)** Bar plot representing million aligned reads per sample, colored by group of samples. **(B)** Heatmap of measured distance between samples, calculated by Manhattan method. Samples have been hierarchically clustered by Euclidean distance. **(C)** Heatmap vst normalized gene expression of top 1000 most variable expressed genes across samples. Gene expression level has been scaled as a z-score, and genes and samples have been hierarchically clustered by Euclidean distance. **(D)** Box plot of estimated fraction of ovary cell types, showing significant differences calculated by Wilcoxon test between groups of samples. In A, B and C, CX_Yh represents Sample X in control group at Y hours; TX_Yh represents Sample X in Treated group at Y hours.

**Supplementary Figure 2.**
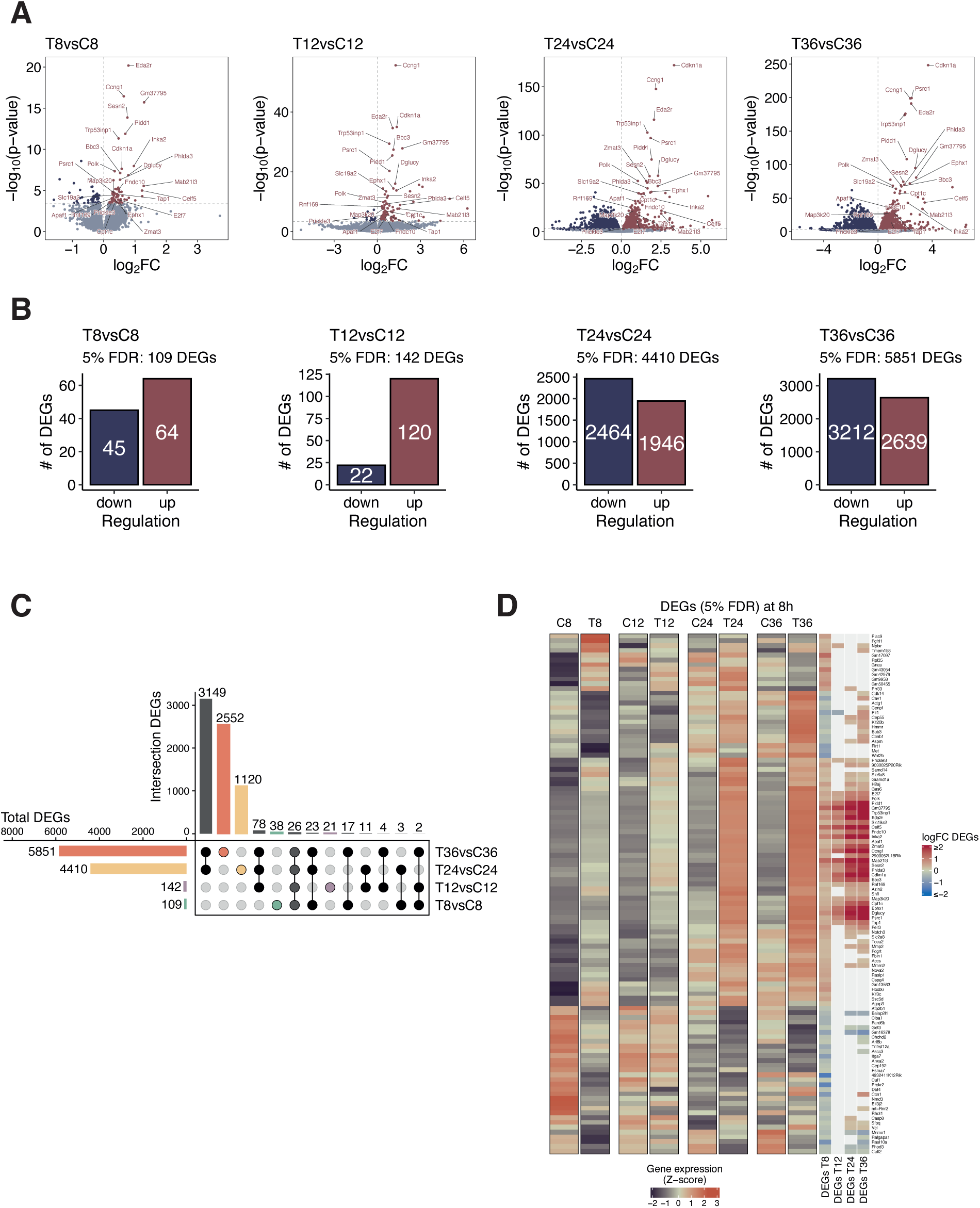
Differential gene expression analyses between CPA-treated and control ovarian samples per time point. **(A)** Volcano plots showing the differential expression levels of genes between CPA-treated and control groups in each time point (8, 12, 24, 36 h). Upregulated DEGs (adjusted p-value < 0.05) are represented in dark-red and downregulated DEGs are in dark-purple. 26 DEGs common to all 4 contrasts are labelled. **(B)** Bar plot of the total number of DEGs per comparison, differentiating between upregulated DEGs (dark red) and downregulated DEGs (dark purple). **(C)** Upset plot of DEGs (adjusted p-value < 0.05) across comparisons from the DGE analyses. Colored points represent DEGs unique to each comparison, black points represent DEGs shared between comparisons, and grey points represent DEGs shared between all comparisons. Left bar plot shows the total number of DEGs per comparison, and top bar plot shows the number of DEGs in each intersection. All possible intersections are shown. **(D)** Heatmap of mean vst normalized gene expression per group of all the DEGs detected at 8 h between treated and control samples (109). Logarithmic Fold Change of the DEGs per time point comparison is indicated at the right, color values are saturated at logFC ≥ 2 and ≤ −2.

**Supplementary Figure 3.**
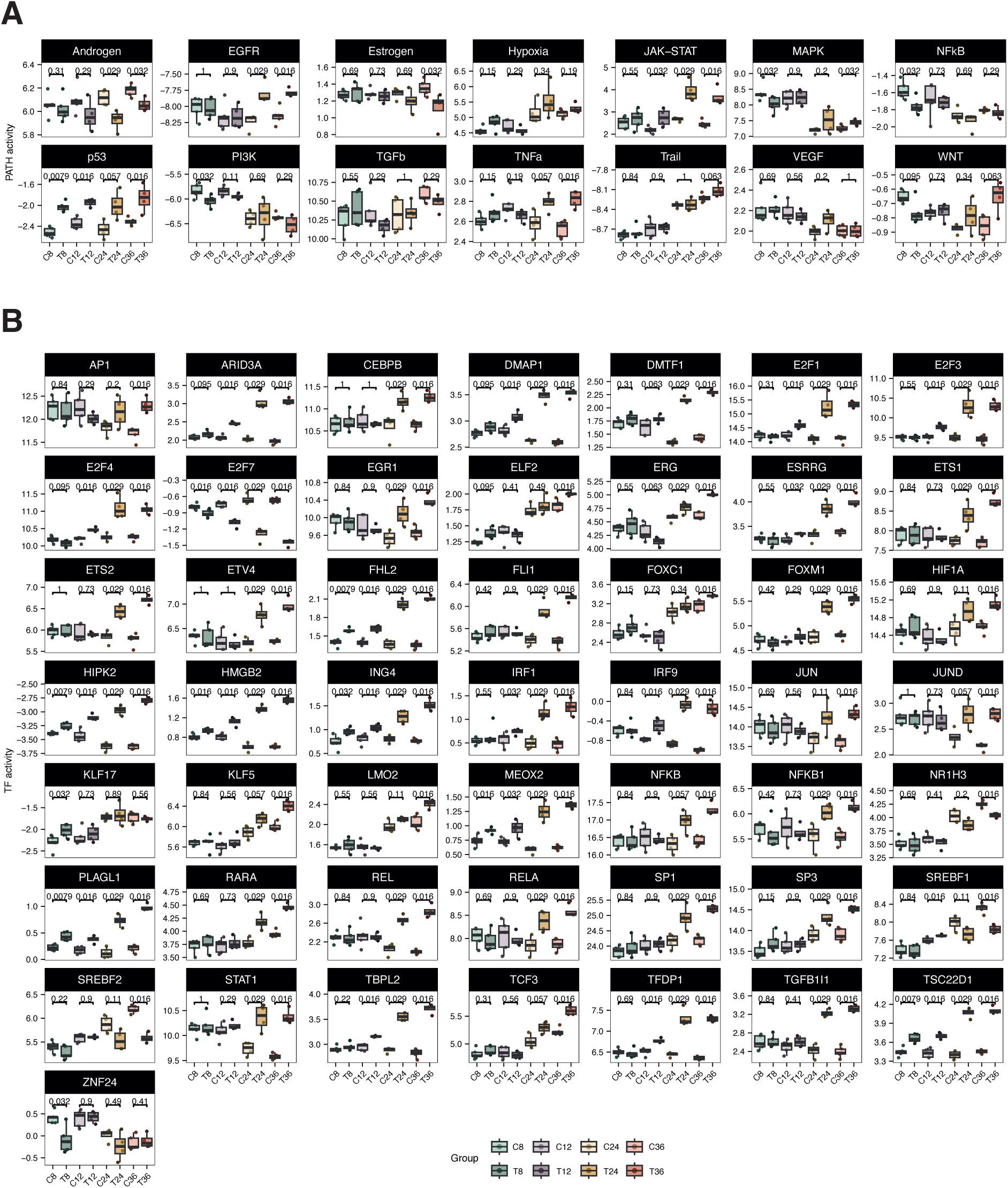
Pathway and TF activity analyses indicating differences between conditions per time point. **(A)** Box plot of pathway activities estimated from the expression of the main 500 pathway-related genes retrieved from the PROGENy database. Significant differences were calculated by Wilcoxon test between groups of samples. **(B)** Box plot of TF activities from the 40 most variable TFs across samples, estimated from targets’ gene expression using CollecTRI database curated regulons. Significant differences were calculated by Wilcoxon test between groups of samples.

## Notes

### Competing Interest Statement

The authors have declared no competing interest.

